# Representational drift without synaptic plasticity

**DOI:** 10.1101/2025.07.23.666352

**Authors:** Caroline Haimerl, Christian Machens

## Abstract

Neural computations support stable behavior despite relying on many dynamically changing biological processes. One such process is representational drift (RD), in which neurons’ responses change over the timescale of minutes to weeks, while perception and behavior remain unchanged. Generally, RD is believed to be caused by changes in synaptic weights, which alter individual neurons’ tuning properties. Since these changes alter the population readout, they require adaptation of downstream areas to maintain stable function, a costly and non-local problem. Here we propose that much of the observed drift phenomena can be explained by a simpler mechanism: changes in the excitability of cells without changes in synaptic weights. We show that such excitability changes can change the apparent tuning of neurons without requiring adaptation of population readouts in downstream areas. We use spike coding networks (SCN) to show that the extent of these tuning shifts matches experimentally observed changes. Moreover, specific decoders trained on one excitability setting perform poorly on others, while a general decoder can perform close to optimal across excitability changes if trained across many days. Our work proposes a simple mechanism without synaptic plasticity that explains experimentally observed RD, while downstream decoding and, by extension, behavior remain stable.

## 1 Introduction

Neurons in sensory areas respond to variations in environmental stimuli with systematic changes in activity. The basic stimulus response of sensory neurons has long been considered fundamentally stable and only transiently modulated given intrinsic dynamics and state changes. This stability has been the focus of studies in recent years as experimental technological advances allowed large numbers of neurons to be tested repeatedly over long time scales, while experimental settings and behavior remained stable. While representational stability has been confimed in some studies [1], others found so-called “representational drift” (RD). Specifically, neurons’ response properties have been reported to change in a seemingly random manner over the time scale of hours, days, weeks and months [2–4]. These RD phenomena have been defined as activity that changes over longer time scales given stable sensation, cognition and action [5]. RD has been described in different brain areas, from sensory to higher-order cognitive and motor cortex. It has been observed with different experimental methods, such as electrophysiology and calcium imaging, and on different time scales, spanning hours, days and months [6–9]. Unlike learning-induced changes in neural representations which are purposeful, highly structured and with a direct impact on behavior, RD appears mostly unstructured and its functional purpose is a topic of debate [10, 11].

Which mechanisms drive RD in these different areas and contexts is unclear. A natural candidate is synaptic plasticity. Synaptic plasticity can produce changes in a neuron’s activity, and several different learning rules have been proposed and tested in the literature to explain the drift observations [12, 13, 10]. Given the stable behavior reported in the RD literature, if one neuron’s activity changes due to synaptic plasticity,these changes need to be compensated by other neuron’s activity to avoid altering overall representations and perception [14, 15]. However, with trillions of synapses in the brain [16] the global coordination of these high-dimensional changes is a nontrivial problem, especially considering that the brain is largely constrained to local learning rules. While previous work has shown how synaptic plasticity could work to update representations in local circuits [17, 10, 18], it is less clear how such mechanisms would deal with the problem of error accumulation over stages of processing. Indeed, given hierarchical sensory processing, any errors accrued at one stage could propagate and amplify in subsequent stages, making the system particularly sensitive to uncompensated changes.

Here we propose that a different mechanism, namely drifts in excitability, may be causing the observed representational drift without impacting downstream readout and consequently keeping behavior stable. Changes in excitability are known to dynamically change activity levels in cells [19–22], resulting from intrinsic processes such as homeostatic regulation [21], fluctuation in intracellular protein levels [22, 23] as well as modulation and input from other brain areas. These fluctuations can occur at similar time scales as the RD observations [20]. Here we show that assuming recurrent connectivity in a population, these excitability changes can also impact metrics such as preferred tuning, rate similarity and trained decoder performance, and are sufficient to explain experimental observations [9, 3, 8, 17]. Importantly, individual synaptic weights do not need to change to achieve drift, suggesting that drift may occur in individual cells without impacting the population decoding at any hierarchical stage, thereby leaving behavior stable.

Our theory and simulation results offer a new perspective on RD with qualitative and quantitative predictions which we confirm in published experimental data. Furthermore, we show how to differentiate experimentally between RD caused by excitability changes versus RD caused by synaptic weight changes.

## 2 Results

Representational Drift has been studied in different brain areas, experimental tasks and using several quantitative metrics. First, RD has been quantified as changes in firing rate similarity, for instance, rate response correlations of cells decrease over days in olfactory cortex [8] and in mouse visual cortex [12]. Second, changes in tuning preferences (e.g. angle) have been reported, for instance, during directed arm movements in macaque motor cortex across different sessions [9]. Third, a widely reported feature of drift is the fading in and out of neurons, quantified for instance in parietal cortex of navigating mice [3, 17], in hippocampus [2, 24], in auditory regions [6, 25], and in several visual cortical regions [19, 7, 26]. Fourth, another hallmark of drift are variations in task or stimulus decodability over days using a fixed decoder fit to a population of neurons to one day, reported for example in auditory cortex [17].

These observations share characteristics such as increased changes over time and functional (behavioral) stability: various measures of behavior do not change given fixed experimentally controlled tasks or sensory stimulations. They also reflect different perspectives on neural coding. When measuring the tuning properties of one neuron we implicitly take a single cell encoding perspective, while population rate similarity or stimulus decodability considers population readout. Here we propose a mechanism that captures these phenomena jointly, while also providing an explanation for how RD leaves function and behavior unchanged.

Let us first consider a simple single neuron model, where a neuron is characterized by its feedforward synaptic weights and its excitability (Fig. 1 A). It receives input reflecting the environment, for instance an image, and its response varies with changes in the environment, such as rotations, which can be summarized in a tuning curve, measuring mean responses over stimulus variations. Under this perspective, any changes in the neuron’s responses must be due to changes in the synaptic (“encoding”) weights (red in Fig. 1 A) or changes in the excitability (green in Fig. 1 A). Specifically, the single-neuron perspective suggests that synaptic weight changes can lead to arbitrary changes in the tuning curve, while excitability-driven changes are restricted to changes in the response gain, or amplitude of the neuron.

**Figure 1.**
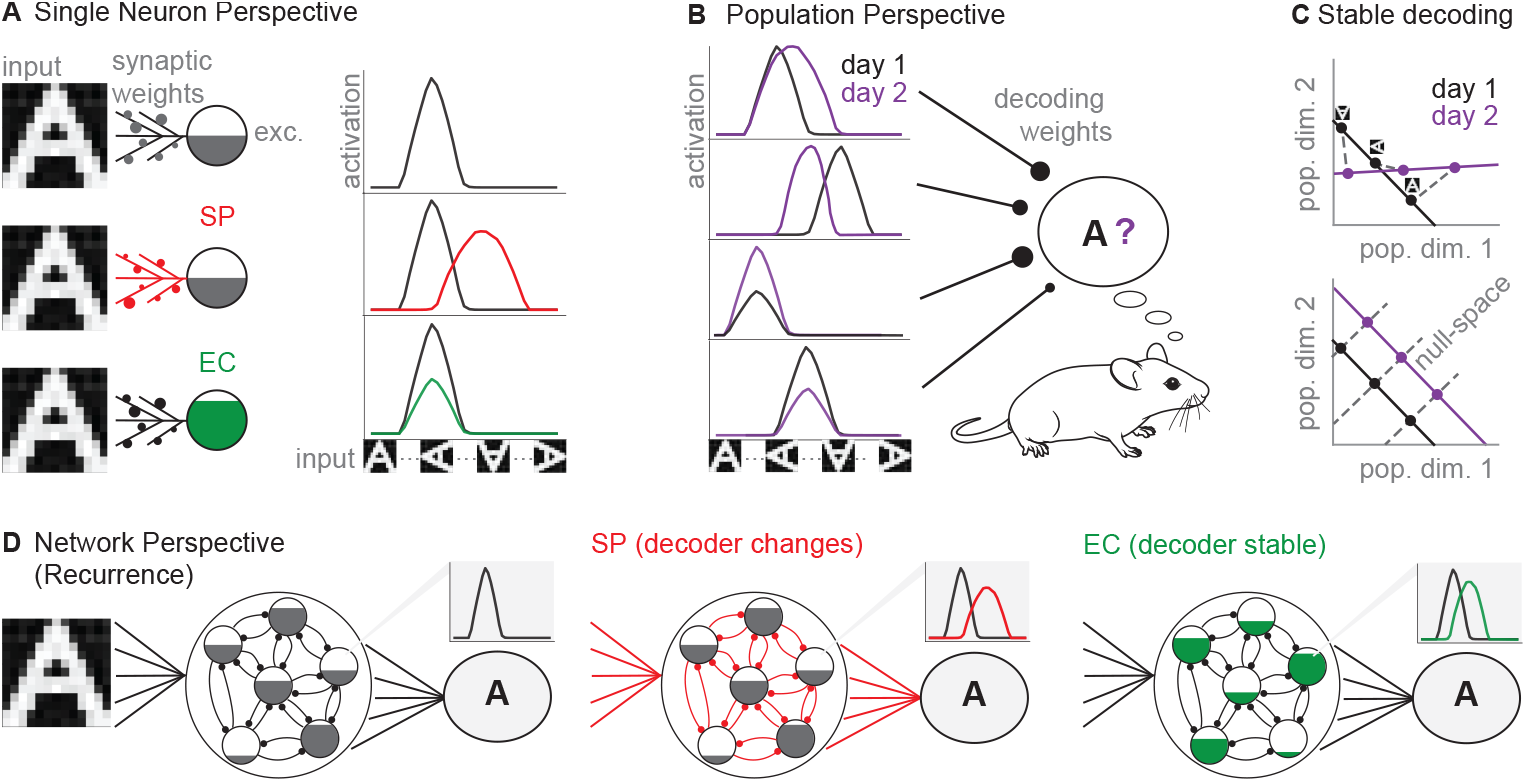
Three different perspectives on representational drift (RD). A) Single neuron encoding perspective. Upper row: A neuron receives input that reflects an environmental stimulus (input). Its response depends on its synaptic weights, and its excitability (exc.). Right: Neurons’ mean response to rotations of the original stimulus (“tuning curves”) can be characterized by tuning preference (position of peak firing), amplitude (peak height of curve), and tuning specificity (width of curve). Middle row: Synaptic plasticity (SP, red) can lead to changes in tuning preference, tuning specificity, and amplitude. Bottom row: Excitability changes (EC, green) only modify the amplitude of the tuning curve. B) Population decoding perspective. A downstream area weights and sums the activity from differently tuned neurons to gain information about the environment and ultimately inform behavior. If tuning of neurons changes due to synaptic weight changes and/or excitability changes (purple), the readout may change, altering behavior. C) To keep decoding stable, there are two possibilities. Top: Activity changes given RD could lead to arbitrary changes in state space. In this case, the linear decoder will need to adjust across days. Bottom: Activity changes given RD could shift along the nullspace axis of the linear decoder. In this case, the linear decoder can remain the same over days. (In both plots, dots indicate similar conditions, e.g. similar images). D) Recurrent network perspective. Left: In recurrent circuits, the tuning of neurons reflects both environmental stimuli and activity of its neighbors. Middle: Synaptic plasticity changes (red) both feedforward and recurrent weights, and also requires changes in downstream decoders to keep the readout stable. Right: Changes in excitability (green) alter only the recurrent input from neighboring neurons, which may change tuning curves of individual neurons. When recurrent dynamics act compensatory, then the readout remains stable.

From the point of view of a downstream area, the stimulus signal is carried by an entire population (Fig. 1 B). A common approximation is that each neuron’s activity is weighted by a set of synaptic (“decoding”) weights which determine its impact on the readout [27]. This readout provides the basis for an animal’s decisions and actions. Therefore, any changes to the neurons’ response properties (reflected in tuning curves) may alter the readout and consequently behavior. In order to keep readout and behavior stable, there are two possible compensatory solutions. First, the response changes could be counterbalanced by a change in the decoding weights [17, 10], see Fig. 1 C, top panel. Second, response changes of individual neurons could be coordinated on the population level such that they are constrained to a so-called ‘null space’ which produces equivalent solutions in decoding space [14, 15], see Fig. 1C, bottom panel. In this case, there would exist general decoders that perform well across several days (although they may be difficult to find in practice, see section 2.3 and Method section 4.4).

Given the slow time scales of representational drift, the only network mechanism for either possibility would seem to be high-dimensional and globally orchestrated synaptic plasticity [10, 17], see Fig. 1 D, middle panel. However, here we demonstrate that slow variations in neural excitability alone can naturally account for the second case, i.e., drift in the nullspace of a representation. Surprisingly, changes in excitability not only cause changes in the gain of individual neurons, but also changes in their tuning preference. These changes occur quite naturally when recurrent connections generate compensatory dynamics across the network (Fig. 1 D). In the following sections, we will show how changes in excitability can explain single neuron and population encoding drift, and further that such changes predict a range constraint for tuning curve shifts, in contrast to models based on synaptic plasticity.

### 2.1 Excitability impacts tuning properties without altering readout

We first illustrate the effect of excitability in a simple spiking network of *N* = 6 integrate-and-fire neurons (Fig. 2 A). The network’s task is to track a two-dimensional sensory input, **x**(*t*) = (*x*_1_(*t*), *x*_2_(*t*)). Concretely, we ask that the input signals can be reconstructed from the spike trains through a linear readout, **y**(*t*) = (*y*_1_(*t*), *y*_2_(*t*)). This readout is generated by filtering the spike trains, weighting them by a set of decoding weights (or coding angles), and then summing. The synaptic conductivities are set to allow the network to perform its task using an efficient spike code, as determined by the spike coding framework (see Methods section 4.1) [28, 29].

**Figure 2.**
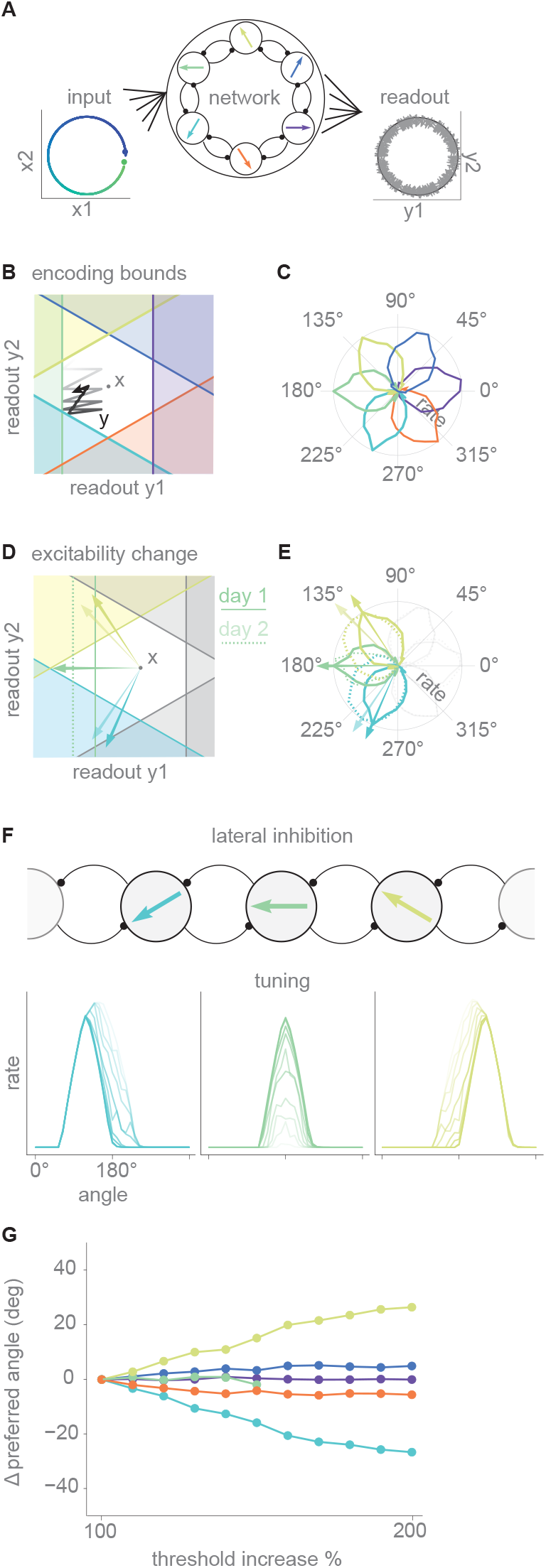
Excitability can impact tuning properties through lateral inhibition. A) A toy model of *N* = 6 neurons with lateral inhibition receives a two dimensional input (left panel). Green to blue gradient indicates time and input angle. Colored arrows within neurons show each neuron’s coding angle (central panel). The population rate vector allows to decode a reconstruction of the input (readout, right panel). B) Network dynamics in readout space. Color-coded lines indicate thresholds of different neurons. White space indicates readout values for which all neurons are subthreshold (“bounding box”). An example readout trajectory is shown in grey-black. Note that the readout cannot leave the white space and thereby remains close to the input x. C) Normalized responses of the six neurons in polar coordinates, given the input shown in A. D) Excitability decreases in one neuron (solid green threshold, day 1; dotted green threshold, day 2). The preferred tuning of the green neuron and its neighbors (yellow, blue) are indicated by arrows (solid arrows, day 1, low contrast arrows, day 2). Note that the preferred tuning of the yellow and blue neurons shifts towards the neuron whose excitability decreased. E) Normalized response of the three colored neurons in panel C before and after the change in excitability (solid lines, day1; dotted lines, day2). Arrows indicate preferred tuning direction before and after. F) Gradual change in tuning curves as excitability of the green neuron decreases. Solid colors, day 1; faded out colors, subsequent days. Note that the tuning curves of the neighboring neurons broaden and shift towards the preferred tuning (180 degrees) of the faded out neuron. G) Change in preferred tuning direction of all neurons as a function of the excitability decrease of the green neuron (% of increase in original threshold). Note that the green line disappears as the activity of the green neurons decreases and tuning can no longer be reliably tested.

Each neuron in the network then receives a weighted combination of the two input signals, as well as lateral inhibition from its neighboring neurons. The voltage thresholds of the six individual neurons can be visualized in the readout space, where they form six different threshold lines, see Fig. 2 B. These lines form a hexagon (white space in Fig. 2 B), which delineates the region in readout space where all neurons are subthreshold. This region bounds the error, (**x** *−* **y**), that is tolerated by the individual neurons before spiking. We will call it the “bounding box” [28].

The dynamics of the network can now be visualized as follows: the readout, **y**(*t*), leaks towards zero in the absence of spiking. When one of the neural thresholds is reached, the respective neuron spikes and pushes the readout back into the bounding box, thereby guaranteeing that the readout stays close to the input signals (Fig. 2 B, grey-black trajectory). Depending on the inputs, the position of the bounding box changes, and different neurons become responsible for keeping the error in check. In Fig. 2 C, we plot the tuning curves of the neurons as a function of the input in polar coordinates.

This picture allows us to jointly consider the effects of synaptic weights and excitability for the example network. The synaptic weights determine the orientation of the neural thresholds, whereas the excitability of each neuron determines the position of its threshold (see Methods section 4.1). The orientation influences for which stimuli the neuron fires, thereby shaping the tuning curve. Less obviously, the excitability also influences neural tuning. When the excitability of one neuron is decreased (green neuron in Fig. 2 D), its gain or maximum firing rate decreases (Fig. 2 E), just as in the single neuron perspective (Fig. 1 A). This results in decreased lateral inhibition of its neighbors, causing the peak firing direction of neighboring neurons to shift towards the coding angle of the inhibited neuron (blue and yellow neurons in Fig. 2 E) and producing a visible shift in tuning curves.

Fig. 2 F,G show what happens if the excitability of one of the neurons further decreases up to the point where it is no longer able to fire. While the respective neuron drops out of the network and its tuning curve disappears, the neighboring neurons’ tuning curves shift their preferred tuning increasingly towards the missing neuron. Biophysically, neighboring neurons experience a decrease in lateral inhibition of the neuron whose excitability decreased, and this decrease shifts their tuning preferences. While the error bounding box deforms slightly, tolerating slightly larger errors in the coding direction of the missing neuron, the input signal keeps being tracked, and the network remains functional.

This example network thereby illustrates two of the key effects of representational drift. First, it captures the fade-in and fade-out of neurons described in experimental observations, where neurons that actively code for an environment on one day no longer do so on a consecutive day, or, vice versa, neurons that are not coding on one day start to do so later [3]. Second, it captures the shift in tuning preferences observed in experiments, which here simply reflects that any decrease in inhibition from fading-out neurons is itself stimulus tuned. Indeed, the recurrent dynamics thereby compensate for changes in the activity of any of the network’s neurons, such that the overall functionality (here, tracking the input) is preserved.

If we assume that the excitability of all neurons in the population smoothly fluctuates over time around some average value, the bounding box itself fluctuates, being slightly more tolerant to some errors on one day and other errors on another day. In turn, individual neurons fade in and out and the preferred tuning of their respective neighbors fluctuate as well. Although these arguments were here made for a simple 6-neuron network tracking a two-dimensional input, we will now show that similar arguments can be made for larger networks tracking multiple input signals. We can thereby test whether the extent of changes obtained through excitability drift can explain experimentally observed and quantified representational drift.

### 2.2 Excitability changes also impact tuning in a high-dimensional spike coding network

We simulate experimental results in a larger network model, where we use a spike coding network with *N* = 2592 neurons (see Methods section 4.1). Instead of a two-dimensional input, we provide a 169-dimensional input (Fig. 3 A). We determine the individual neuron’s coding angles by choosing Gabor filters as a concrete example [30], see Fig. 3 B. These filters are common models of neural response profiles in primary visual area, V1. We emphasize, however, that we envision this model as a generic example of high-dimensional computations in neural circuits. Indeed, the results below do not depend on the choice of Gabor filters, but are universal features of spike coding networks with redundancy. The key advantage of the Gabor filter example is that it allow us to measure neural tuning through angles, just as in the two-dimensional case. It is straightforward to construct other SCNs with angular variables, e.g., for the head-direction system or for entorhinal cortex [31].

**Figure 3.**
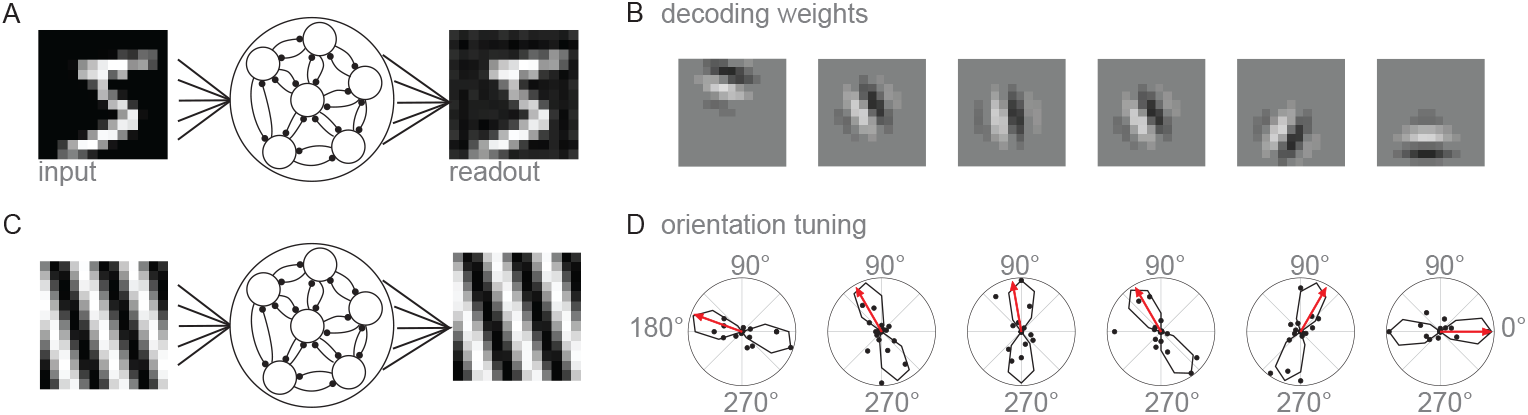
A 13^2^ dimensional pixel space is encoded by a SCN with *N* = 2592 neurons. A) Population encodes image input so that it can be reconstructed from its spike trains. B) Example neurons’ coding weights in image space (arranged in 13*×*13 space). C) Neurons’ tuning is probed by presenting them with drifting gratings of different directions. D) Mean responses rate to the presented drifting gratings of example neurons in B (rate in Hz along radius axis). Dots indicate the mean response. Lines show the bounded sine curve fits. Red arrows indicate coding angles of each example neuron. Neurons’ thresholds are set uniformly for this simulation.

We first note that the network accurately represents its input, as illustrated by the decoding of an image from the neural population (Fig. 3 A). We test the neurons’ orientation tuning by presenting it with 16 drifting grating images and measuring neurons response properties (Fig. 3 C). Because of the recurrent dynamics, not all neurons are active or show strong tuning (despite their Gabor weights), as they inhibit one another. However, we observe significant tuning in *∼* 20% of the population in a given session (see Methods section 4.2 for details). Fig. 3 D shows data and tuning curves fit for example neurons. Note that here neurons’ thresholds are set uniform across the population (no drift), leading to well-aligned coding angles (red arrow) and tuning curves.

We then test the effect of drifting excitability. Even though causes of excitability changes can be diverse, we can summarize them as changes in the threshold of our neural spiking model. We allow neurons’ thresholds to vary slightly over days by adding Gaussian noise for every simulated day, but limiting individual thresholds by a lower bound (see Methods section 4.3 and Fig. 4 A, bottom panel). This allows us to test the effect of excitability changes in a high-dimensional neural population on single-neuron tuning properties and on population coding.

**Figure 4.**
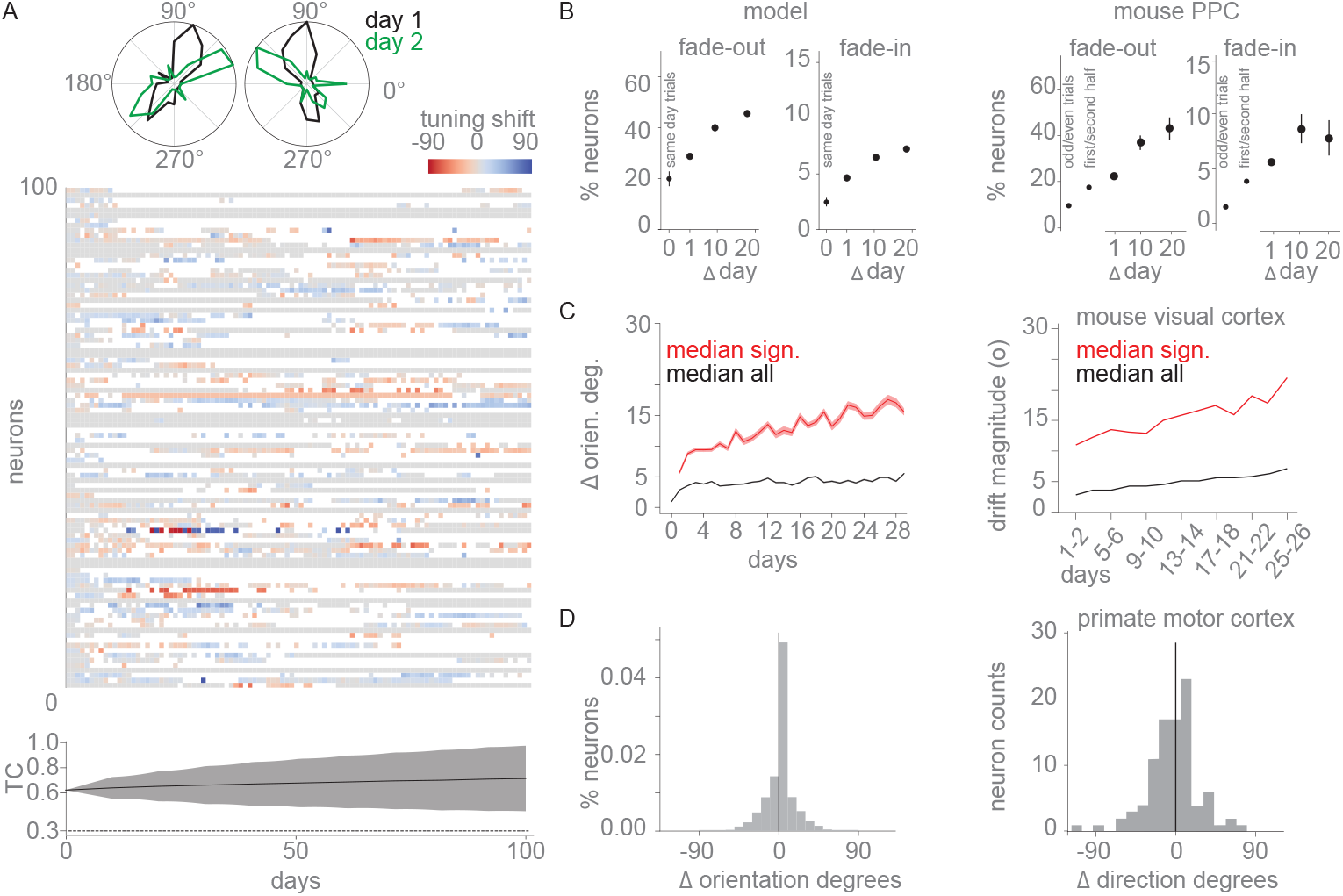
Neurons’ tuning properties vary with random drifts in excitability. A) Top: Tuning of two example neurons on two days, plotted as in Fig. 3D. Middle: Tuning changes over 100 days. Color indicates tuning changes of 100 subsampled neurons (out of the 2592) relative to day zero in degrees. White indicates a neuron has become untuned. Bottom: The distribution of threshold changes (TC) across the population over time (mean and std). Dashed line indicates the lower bound on threshold distribution. B) Left: Percentage of neurons that fade out or fade in after 1, 10 or 20 days in the network model, and cross-validated fade-in/out for different trials of day zero (see Methods section 4.2). Right: Experimental results from mouse posterior parietal cortex (PPC) on fade in/out, replotted from Driscoll et al. [3]. Mean and standard deviations are computed across model instantiations (left) or experimental sessions (right). C) Left: Median and SE of preferred orientation changes over days across populations, for all neurons (black) and only significantly tuned neurons (red). Right: Experimental results from mouse visual cortex, replotted from Bauer et al. [32].D) Left: Distribution of preferred orientation changes over all neurons and days. Right: Experimental results from primate motor cortex on tuning changes over sessions, replotted from Rokni et al. [9].

We simulated the network’s activity when subjected to excitability drift over many days. Fig. 4 A, top panel, shows two example neurons’ tuning curves changing over two example days, and visualizes the changes in preferred tuning of subsampled neurons together with their simulated threshold changes (see Methods section 4.3 & 4.2). We make several observations. First, neurons fade in and out of the population code, as indicated by the many white spots in which the respective neurons are no longer tuned. Second, many of the neurons change their preferred tuning over time. Third, we observed that the reconstruction error under the ground truth decoder, quantified as the mean squared error (MSE) over all tested drifting grating directions, stays constant (data not shown, but see Fig. 3 A for an example reconstruction). The network thereby preserves its output despite the observed changes in the neural representation, as suggested by the intuitions developed in the small toy model with *N* = 6 neurons above.

### 2.3 Excitability changes can explain experimentally observed changes in single neuron encoding

To compare the network behavior with experimental data, we first studied the fade-in and out of neurons. If a neuron’s excitability decreases strongly relative to its neighbors, its neighbors become disinhibited and take over the coding space, resulting in that neuron’s fade-out, i.e., it no longer actively participates in coding for the stimulus. Such fade-out may be reverted at any time as the relative strength of excitability changes. Given a fixed time (day zero), we can quantify the fraction of neurons that have faded out (or faded in) on a subsequent day, as shown in Fig. 4 B. We find that fade-in and fade-out in simulations with small excitability changes can quantitatively explain the percentage of neurons which stop or start coding for spatial environments in experimental data [3] (Fig. 4 B). Even though the active population changes from day to day, the percentage of actively coding (tuned) neurons stays approximately constant (data not shown).

In a second step, we studied the changes in preferred tuning over time when excitability drifts according to a random noise process (see Methods section 4.3). This is illustrated in greater detail in Fig. 4 C. We quantified the amount of tuning drift by computing the angular difference between neurons’ preferred orientation on each day compared to the first day. We find a smooth increase in changes over days (Fig. 4 C, similar in extent and dynamics to experimental observations from mouse visual cortex [32]. Notably we find that tuning changes are correlated across neurons in a population (see correlated mean changes in Fig. 4 C). Across days we find a distribution of changes that is centered around zero and falls off well before the maximum difference of *±*90 degrees for orientation tuning (Fig. 4D). This distribution matches what has been found experimentally in angular direction tuning of motor cortex [9], although we find more neurons with close to zero changes in tuning (similar to [32]). The falloff of the distribution can be explained by the particular impact that threshold changes have on the population coding geometry. Specifically threshold changes move the coding boundary of one neuron relative to the other neurons, thereby pulling or pushing the other neuron’s preferred tuning orientations (Fig. 2 D,E). However, since they cannot change the weight-fixed coding angle of any neuron, the maximum pull or push of the preferred orientation is limited. Our results therefore propose that angular tuning changes due to representational drift (and without learning) should be limited and vary smoothly over time, eventually converging to a unimodal distribution of changes.

We note that this limit in the distribution of changes arises because of the fixed decoders. When the decoders can change, as in networks based on synaptic plasticity, this constraint does not appear naturally. To illustrate this, we simulated Hebbian learning in the toy SCN (described in section 2.1) with noise on the angular weight updates [33], similar in spirit to [10] (see Supplementary Figure 2 for details). Networks drift between different near-optimal configurations of feedforward and recurrent weights and represent the stimulus well and stably. However, due to the noise term on the weight updates, there is drift between different solutions to the encoding-decoding problem, resulting in drifts in tuning which are not bounded, eventually resulting in a uniform distribution of changes.

### 2.4 Excitability changes can explain experimentally observed changes in population coding

We then consider the impact of representational drift on the population level and test whether individual neurons’ excitability changes can explain both population drift and stable population coding. We find that population activity vectors, made up of the mean rate of each neuron for a specific input, become increasingly dissimilar over days with excitability changes (Fig. 5 A). Specifically, we measure the correlation of population activity vectors on each simulated day compared to the first day and find a decrease in correlation. This decrease is quantitatively similar to what has been observed experimentally in olfactory cortex [8] (see Fig. 5 A) and in hippocampus [10] (not shown).

**Figure 5.**
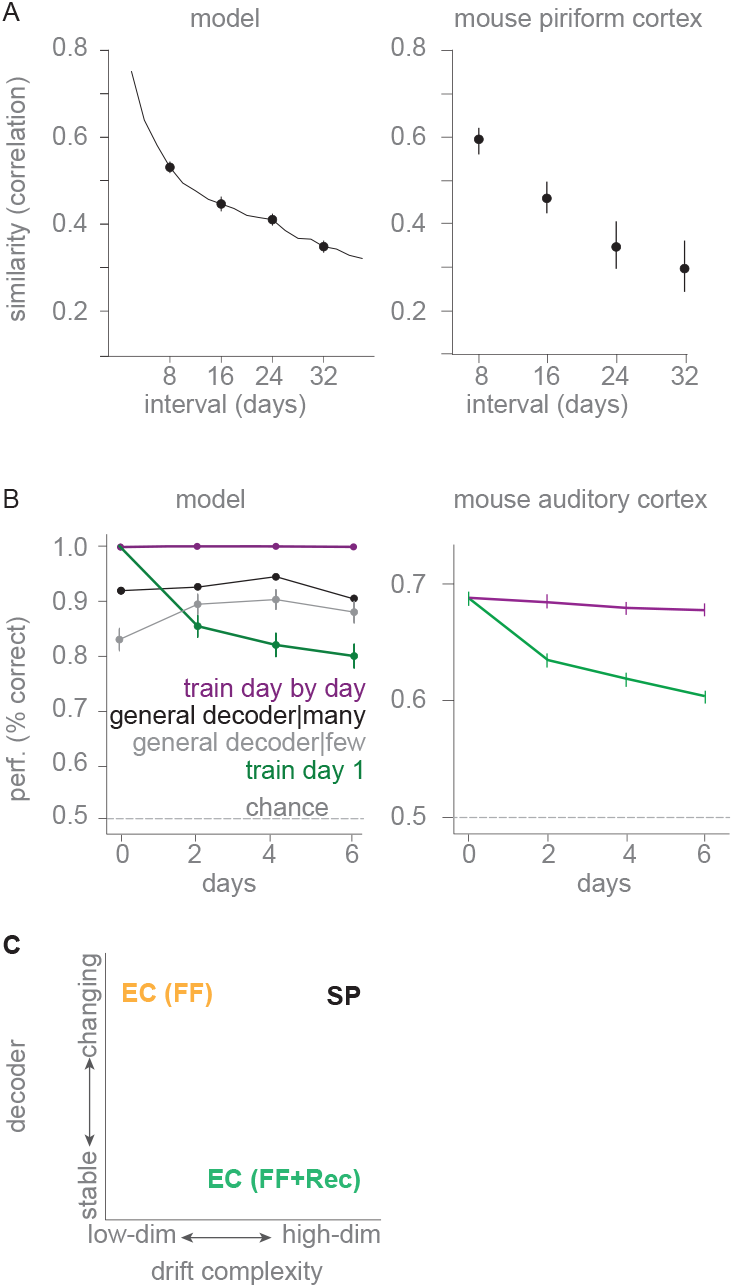
Population encoding drifts smoothly over days. A) Left: Population rate similarity, measured as the correlation between the population activity vector on the first day and subsequent days. Plotted are mean and SE over population and over 10 simulations. Right: Experimental results from mouse piriform cortex responses to odors over 32 days (Schoonover et al. [8]). B) Left: Logistic regression decoding of drifting gratings (Fig. 3C) over time from 5% of neurons in the population. Purple: retrained daily on 7 trials and tested on 3 held out trials of the same day. Green: trained once on day zero on all 10 trials and tested on subsequent days. Black & grey show general decoders that are trained on 10 trials sampled across days except the one that testing is performed on. Black shows performance if 10 training trials were drawn from all simulations except test day (99 out of 100 sessions with 10 trials each). Grey shows performance if training trials were drawn from only three sessions. Mean and standard error are shown for 100 decoding simulations. Right: Ex-perimental results from mouse auditory cortex (As-chauer et al. [6]). C) Hypotheses such as excitability changes (EC) and synaptic plasticity (SP) that may cause representational drift but preserve behavior.

RD is characterized by stable experimental behavioral performance [5], suggesting that downstream readout is not affected by the tuning drifts in a given encoding population. However, if a logistic decoder is fitted to neural activity recorded on a particular day, its performance drops smoothly but substantially across subsequent days [6]. Changes in excitability offer an explanation for why observed decoding performance may decrease, while neural readout as used by the brain stays stable, leading to persistent behavioral performance in experimental tasks. Specifically, excitability does not change the actual coding angle of a neuron, that is, the orientation of the bound in readout space (see Fig. 2 C). Instead, the overall geometry of the space changes such that neurons fire maximally to orientations that deviate from their actual coding angles. These changes occur because neurons’ responses are heavily influenced by the recurrent dynamics of the network. However, statistical fits of linear decoders will be steered by preferred tuning rather than the (hidden) actual coding orientations, thereby leading to misestimates of linear decoders.

We illustrate this theoretical intuition by simulating decoding from our population in two ways. We aim to discriminate pairs of drifting grating directions (Fig. 3 C) by fitting a logistic regression model to a subsample of neural responses, given that experimental datasets usually include only a subsample of the total population of coding neurons. We measure the percentage of correctly identified drifting gratings to evaluate the logistic regression (see details in Methods section 4.4). The optimal model is one that is trained and tested on simulated data from the same day and therefore best adapted to the coding geometry of this day’s excitability settings (purple in Fig. 5 B). We compare this “upper bound” of performance to a decoder that is trained only on the first day and then tested on subsequent days without readjustments of the weights (green in Fig. 5 B). We find that decoding performance of such a single day decoder drops substantially if excitability changes, similar to what has been observed in experimental data of categorizing sound from auditory cortical neurons [6] (replotted for comparison in Fig. 5 B). For the specific network simulated here, we find that a general decoder trained on data across different days can recover much of the optimal decoder’s performance, suggesting that the ground truth stable decoder can be recovered with sufficiently heterogeneous data (black and grey lines in Fig. 5 B). This suggests that as experimental methods allow tracking the same neural populations over increasingly long-time scale, fitting general decoders over sufficiently many days will provide better estimates of the true optimal decoders for these populations.

To summarize, possible drift mechanisms can be characterized along two dimensions. First, the extent of tuning changes they permit, which we term “drift complexity”. For instance, amplitude changes of tuning curves are low complexity, while arbitrary shape changes of tuning curves are high complexity. Second, the extent of changes required to assure robust behavioral performance. Large changes are required if a downstream decoder needs to be adapted on a constant basis, and no changes are required if downstream decoding remains stable (Fig. 5 C). Given these two axes, we can distinguish three possible RD mechanisms. Synaptic plasticity (SP) can explain (almost) arbitrary tuning curve changes (high drift complexity), but requires adaptation of downstream decoders. Excitability changes in a feedforward circuit can only produce simple drifts in amplitude (low drift complexity) and also require adaptation of decoders. Excitability changes in a recurrent circuit, however, causes changes in tuning preferences as well as amplitude (intermediate drift complexity), and, when recurrent activity is compensatory as in the spike-coding network, can work with stable decoders. Here, we showed that the intermediate drift complexity explained by excitability changes is largely compatible with the drift complexity observed in experimental work. Excitability may therefore be sufficient to explain neural and behavioral hallmarks of RD.

## 3 Discussion

Our study proposes a simple mechanism explaining the experimental phenomena of representational drift in neural populations. We demonstrate that changes in neuronal excitability of recurrently connected populations can lead to drift in neural representations. This drift manifests itself in the dropping in and out of neurons and in changes of tuning preferences. We show that these changes can be dynamically coordinated through the recurrent neural activity such that a stable population code is maintained.

In past work, representational drift has often been attributed to ongoing synaptic plasticity. Indeed, representational drift can arise from noise in synaptic updates within a degenerate solution space [10]. Such a mechanism puts few limits on the type of changes in neural tuning that should be observable. As shown in Supplementary Fig. A.1, e.g., neurons should be able to completely reverse their tuning. This prediction is quite different from ours, where neurons angular tuning is always anchored and tuning changes follow a normal distribution with a more limited standard deviation.

An alternative suggestion is that drift arises from behavioral variability, i.e., because experimental conditions change over days, and those changes are not accounted for. This mechanism was emphasized by Sadeh and Clopath (2022) analyzing visual recordings from mouse. Furthermore, in hippocampal recordings in freely flying bats, ‘drift’ in representations could be fully attributed to systematic changes in their flying path [34]. Our model is compatible with these findings, as a change in an (undetected) input to a neuron causes an effective change in the neuron’s excitability.

Here, we propose excitability as the key mechanism underlying representational drift. Besides changes in (unobserved) inputs into neurons, changes in excitability may be caused by a wide range of processes with different time scales and magnitude. For example, short-term rapid (sec-min) changes in intrinsic excitability can be produced by phosphorylation of ion channels [35, 36], while synaptic scaling operates over longer timescales (hours to days) as it involves changes in the expression or trafficking of synaptic proteins [37]. Even longer timescales of excitability changes have been reported by the effects of the brain-derived neurotrophic factor which can sustain day-long changes [38]. Many of these processes are believed to support homeostatic regulation at the level of single cells or populations. This mechanism is consistent with various normative explanations of representational drift, such as its role in regularization or exploration of different solutions [10, 11]. Importantly, our proposed mechanism is not limited to the visual system. The combination of recurrent connectivity and lateral inhibition, which are ubiquitous features of neural circuits, allows this excitability-based drift to occur in various brain areas, consistent with the experimental findings of drift across sensory, cognitive and motor regions. Our framework differs from previous work which suggested that excitability changes produce representational drift by driving changes in synaptic weights [20]. In our work, the excitability changes produce drift by driving changes in the recurrent dynamics rather than the synaptic weights.

Our framework makes several testable predictions. We expect the amount of drift to be constrained by the consistent geometries of the coding space. In the case of angular tuning, studied here in detail, we expect drift to be normally distributed with constrained maximum tuning changes if caused purely by excitability changes, since large tuning curve changes would require synaptic weight changes. This prediction is stricter than the requirement of a decoder null space, and distinguishes our model from those based on synaptic plasticity. The prediction may also be more amenable to experimental tests, as establishing a coding null space from experimental data is fraught with difficulties: it is often impossible to establish the ‘ground truth’ variables that need to be decoded, and one may suffer from biases due to the subsampling of neural populations [39, 40].

In brain areas with extensive mixed selectivity, quantifying tuning shifts may be more challenging due to the complex nature of neuronal responses. Here, representational similarity has been established as a common quantification of drift [19, 12, 24, 32]. We replicate these results in Fig. 5 A, showing that excitability alone can reproduce the results of decreasing representational similarity.

Our model is based on a spike-coding network that reproduces its input [28, 29]. This type of autoencoder network has been studied before in the context of neuron loss [41, 28]. Neuron loss can be seen as an extreme case of a change in excitability, in which a neuron stops firing altogether. Our network replicates these previous findings in such cases. While we here focused on networks that reproduce their input, we note that this specific computation is neither a core assumption nor limiting factor of the underlying model. The only component of this model necessary to obtain representational drift is a boundary made up of individual neurons (Fig. 2). This will generally be the case if the connectivity of a network is low-rank and effective thresholds are properly set [42]. Many other types of computations besides auto-encoding can be performed by such networks [43, 42]. As long as functionally neighboring neurons perform similar computations, RD will occur in these networks as well when excitabilities fluctuate over days.

In conclusion, our study offers a unifying framework for understanding representational drift across different brain regions. By highlighting the role of neuronal excitability changes and homeostatic plasticity, we provide a mechanistic explanation for the coordinated nature of drift observed in neural populations. This work not only reconciles seemingly contradictory observations but also generates testable predictions for future research,paving the way for a deeper understanding of the dynamic nature of neural representations.

## 4 Methods

### 4.1 Spike Coding Networks

We use the spike-coding network (SCN) framework to model how neurons’ activity is shaped by their feedforward input weights **F** and recurrent connectivity **W** [29, 28, 42]. As a concrete instantiation, we chose a network that tracks a multivariate input [28] and is dominated by lateral inhibition. Given a fixed matrix of readout weights, **D**, we choose feedforward weights, **F** = **D**^*T*^, and recurrent connections, **W** = *−***D**^*T*^ **D**. For simplicity we set the columns of **D**, called **D**_*n*_, to have unit norm. We refer to **D**_*n*_ as the decoding vector of neuron *n*, which determines the neuron’s coding angle.

Given a set of signals, **x**(*t*) = (*x*_1_(*t*), …, *x*_*M*_ (*t*)), the neurons are fed the inputs (*k* = 1 … *M*),

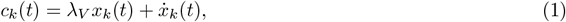

so that the voltage of the *n*-th neuron evolves as,

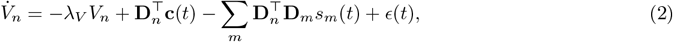

where *s*_*m*_(*t*) is a series of delta-functions that model the spike trains, *λ*_*V*_ is the filtering time constant and *ϵ* is gaussian-distributed time varying voltage noise with 0-mean and *σ*_*ϵ*_ variance (see Table 1 for parameter details). The effective time scale of the neurons is set by *λ*_*V*_ which is set to 100 and can be thought of as 10ms. Spikes are generated whenever a threshold *T*_*n*_ is reached, and the voltage reset is integrated in the connectivity matrix. Accordingly, the neurons are modeled as (current-based) leaky integrate-and-fire neurons. We furthermore define the readout as a filtered summation of the spike trains, weighted by the decoding vectors,

**Table 1.**
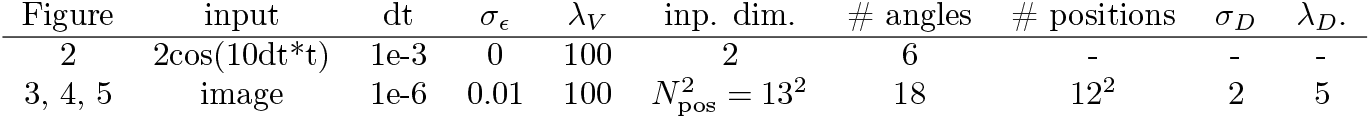
Parameters used for simulations in each figure. The parameters are: *dt*, the time step in sec; *σ*_*ϵ*_, the standard deviation of the voltage noise *ϵ*; *λ*_*V*_, the filtering time constant of the neurons; the dimensionality of the input **x**, the number of angles or spatial receptive field positions along which neurons are distributed; *σ*_*D*_,the standard deviation of the Gaussian envelope of the Gabor filter on the spatial grid defined by 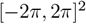 and *λ*_*D*_, the wavelength of the Gabor filter’s sinusoidal factor. Note that *σ*_*ϵ*_ is further scaled by 2 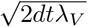 in numerical simulation. Furthermore, the “input dimension” does not include the additional background input, which is independent and regulates the firing rate.

| Figure | input | dt | $\sigma_\epsilon$ | $\lambda_V$ | inp. dim. | # angles | # positions | $\sigma_D$ | $\lambda_D$ |
| --- | --- | --- | --- | --- | --- | --- | --- | --- | --- |
| 2 | $2\cos(10dt*t)$ | 1e-3 | 0 | 100 | 2 | 6 | - | - | - |
| 3, 4, 5 | image | 1e-6 | 0.01 | 100 | $N_{\text{pos}}^2 = 13^2$ | 18 | $12^2$ | 2 | 5 |

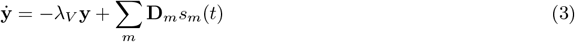

In Fig 2 A&B, we set the coding angles to uniformly tile the two-dimensional input space, and we set all thresholds to a fixed value (*T*_*i*_ = *T*). The relation between a neuron’s voltage and thresholds is then given by [28],

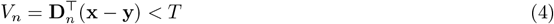

and, when visualized in the space of readouts, gives rise to the bounding box. The input signal in Fig 2 follows a circle in a two-dimensional space, so that **x**(*t*) = (*a* sin(*bt*), *a* cos(*bt*)). In turn, neighboring neurons have similar tuning and tuning curves are equidistributed in angle space. In Fig 2 C-E, one of the neuron’s thresholds is incrementally increased, until it stops firing.

For Fig 3-5, the neurons’ input weights were modeled as local oriented gratings, or Gabors (see Table 1 for parameter details). Each neuron’s input weight filter is defined on a two-dimensional discretized spatial grid with filter center position (*µ*_*i*_, *µ*_*j*_) *∈* [*−*2*π*, 2*π*]^2^ and angle orientation *θ*_*D*_ *∈* [0, 2*π*]. More specifically, we mapped indices of the spatial grid onto the elements of the decoding vector, (*i, j*) *→ k*, so that for the *n*-th decoding vector,

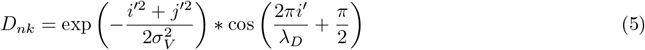

where

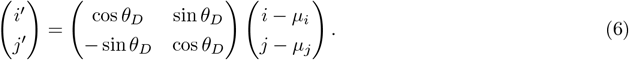

The parameters *σ*_*D*_ and *λ*_*D*_ control the width of the Gaussian envelope and the wavelength respectively. The coordinate system is rotated by *θ*_*D*_ and centered on ***µ*** before applying the cosine modulation. Gabor filters are generated for all combinations of 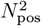 positions and *N*_ang_ angle orientations. Values of the filter with absolute magnitude below 0.1 are thresholded to zero.

Following [41], we furthermore added a constant background signal that can be read out through the overall population firing rate. The additional input signal regulates the base firing rate of the neurons and prevents the generation of seizure-like ping-pong states in these networks (see [28, 41] for detailed discussion). The final coding angles were therefore **D**_*n*_ = (**G**_*n*_, *z*) where **G**_*n*_ is the *n*-th Gabor filter and *z* the constant coding dimension. Thresholds were drawn from a normal distribution with weight-matched mean 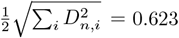 0.623. In numerical simulations of the spiking dynamics, we used the Euler method with the additional rule that only one neuron spikes within the time window set by the step size *dt*. This extra rule allows us to choose a larger step size and thereby speeds up the simulation. The code for simulations is available upon request.

### 4.2 Assessing neurons’ direction and orientation tuning

We quantify tuning of neurons by computing the average spike count for each stimulus angle. For the twodimensional input simulations we binned the stimulus angles in 10degree bins. Where voltage noise was used (simulations in Fig. 4), we compute tuning curves on trial-averages and test on left-out trials. For the image-input simulations, we assessed whether neurons were significantly tuned by fitting a half-wave rectified sine curve to normalized tuning curves (two peaks for assessing orientation tuning, mean rates normalized to lie between 0 and 1) and compared the mean squared error (MSE) of the sine curve fit to that of a constant mean fit. If the MSE of the sine curve fit was lower than the constant mean fit error, we considered them significantly tuned. Among the not-significantly tuned neurons, we tested whether a sine curve with a single peak would improve the fit (direction tuning), but the fraction of neurons whose fit was improved was only*∼* 1% and hence negligible. To assess changes in preferred orientation for significantly tuned neurons, we computed the difference in the phase of sine curves. When we use voltage noise (Fig. 4&5), we fit tuning on 5 simulated trials and test fit significance on another 5 simulated trials. Note that when neurons are orientation tuned the maximum amount of tuning changes cannot be larger than 90 degrees.

### 4.3 Excitability drifts

We model intrinsic and extrinsic factors that cause excitability changes jointly as effective changes in the threshold *T*_*n*_. Accordingly, the value of *T*_*n*_ does not reflect the actual biophysical neural threshold, but rather summarizes physical sources that jointly set a neuron’s excitability. In the high-dimensional input space of Figs. 4&5, we create threshold sequences through two steps which provide us better control over the size of the threshold changes. For one neuron we start with a threshold sampled from a baseline distribution (gaussian with lower limit, parameters see Table 2) and then sample zero-mean gaussian noise to be added to previous thresholds. Second we interpolate between these generated thresholds to create smooth steps of threshold changes. If at any point a threshold drops below 0.3 we set it to 0.3 to avoid neurons drifting too far into the coding space. The exact lower bound threshold is not crucial and results are similar for variations. We provide a detailed list of parameter choices for threshold changes in Table 2.

**Table 2.**
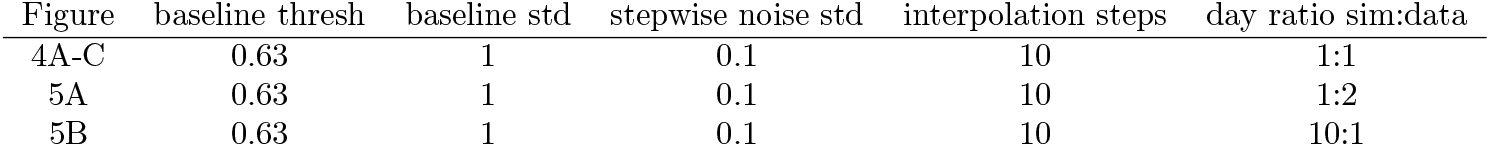
Parameters used for threshold drift simulations in each figure. The baseline threshold defines the mean of the distribution and is set to the norm of the weights. The baseline std defines the standard deviation of the threshold distribution from which thresholds are sampled on the first day. The stepwise noise std defines the standard deviation of the 0-mean noise that is added to the old threshold draw for every new threshold draw. Thresholds can not decrease below 0.3. The interpolation steps defines how many steps are in between an old threshold draw and a new threshold draw, so that in total the simulated threshold trajectories vary smoothly from one threshold draw to another. The ratio sim:day defines how we matched simulated thresholds to experimental days, for instance a ratio of 10:1 would mean that 10 simulation steps would translate to one day in the experiments.

### 4.4 Decoding

For drift direction discrimination, we use logistic regression trained on the population rates to discriminate pairs of drifting gratings, similar to [6]. Most experimental setups do not allow recording from all neurons in a population. In order to take this into account when comparing our decoding results, we decode from only a subsample of neurons in the population. We report performance as % correctly discriminated direction pairs, averaged over all pairs (as in [6]) and over 100 times sampling 5% of all simulated spiking neurons. Each time we use 7 trials for training and 3 for testing (unless training and testing on different days). Relative performances are similar for different percentages of neuron subsampling although absolute performances change.

### 4.5 Synaptic weight changes lead to arbitrarily large changes in tuning

In our simulation of networks encoding two-dimensional input spaces, we assumed optimal weights *D*. We can instead learn these weights by allowing plasticity of feedforward weights *F* and recurrent weights Ω according to a simple Hebbian update rule, where weights are updated whenever a neuron spikes, according to [33]:

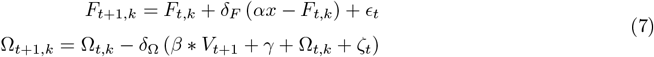

where *δ*_Ω_ = 0.001 and *δ*_*F*_ = 0.0001, *α* = 0.18 and *β* = 1.39. The learned weights allow for a good reconstruction of inputs and converge to optimal solutions (for details see [33]). However, there exists an infinite space of optimal solutions that may be learned, which can be thought of as arbitrary rotations of the bounding box (Supplementary Fig. A.1 A, middle). Noise added to the weight updates therefore allows the network to traverse this solution space and produce dynamically changing coding geometries in the process (Supplementary Fig. A.1 A, right). These dynamically changing representations do not form optimal bounding boxes at any particular point in time but drift between optimal solutions. Specifically, we can test the effect of noise on the update steps, similar to [10]. We use noise *ϵ*_*t*_ and *ζ*_*t*_ drawn from an Ornstein-Uhlenbeck process with *τ* = 0.05 and *σ* = *δ*_*F*_ *α*0.4 for feedforward weights and *σ* = *δ*_Ω_*α*0.4 for recurrent weights. We compute the coding directions over time and find dynamic variations where individual neurons coding directions drift and neurons repel each other so as to push towards uniform tiling of the space, although under sufficiently large update noise this optimal uniform tiling is not achieved (Supplementary Fig. A.1 B). We compute tuning curves across days for example neurons and show that, in contrast to excitability changes (Fig. 2 D), synaptic weight changes produce unconstrained tuning curve changes (Supplementary Fig. A.1 C). These weight-dependent tuning curve changes eventually span the entire spectrum of preferred direction tuning changes (Supplementary Fig. A.1 D). While the distribution of changes starts out Gaussian, with increasing length of the simulation, changes become increasingly uniform (Supplementary Fig. A.1E). This is in contrast to experimental findings where changes in direction tuning are constrained to *±*90 degrees [9] (see Fig. 4D).

## Acknowledgments

We are especially thankful to William Podlaski and Guillermo Martín-Sánchez for their pivotal ideas during the early stages of this project. We furthermore thank Francesca Mastrogiuseppe, David Lipshutz and Laura Driscoll for helpful discussions and comments on the manuscript and Joel Bauer for sharing his data with us. CH is supported by a Transition to Independence grant from the Simons Foundation Collaboration on the Global Brain (code 00007243). CKM is supported by the Simons Foundation Collaboration on the Global Brain (code 543009 and 2794-04), as well as NIH R01 EY035896 and NIH RF1 NS127107.

## A Supplementary Material

**Figure A1.**
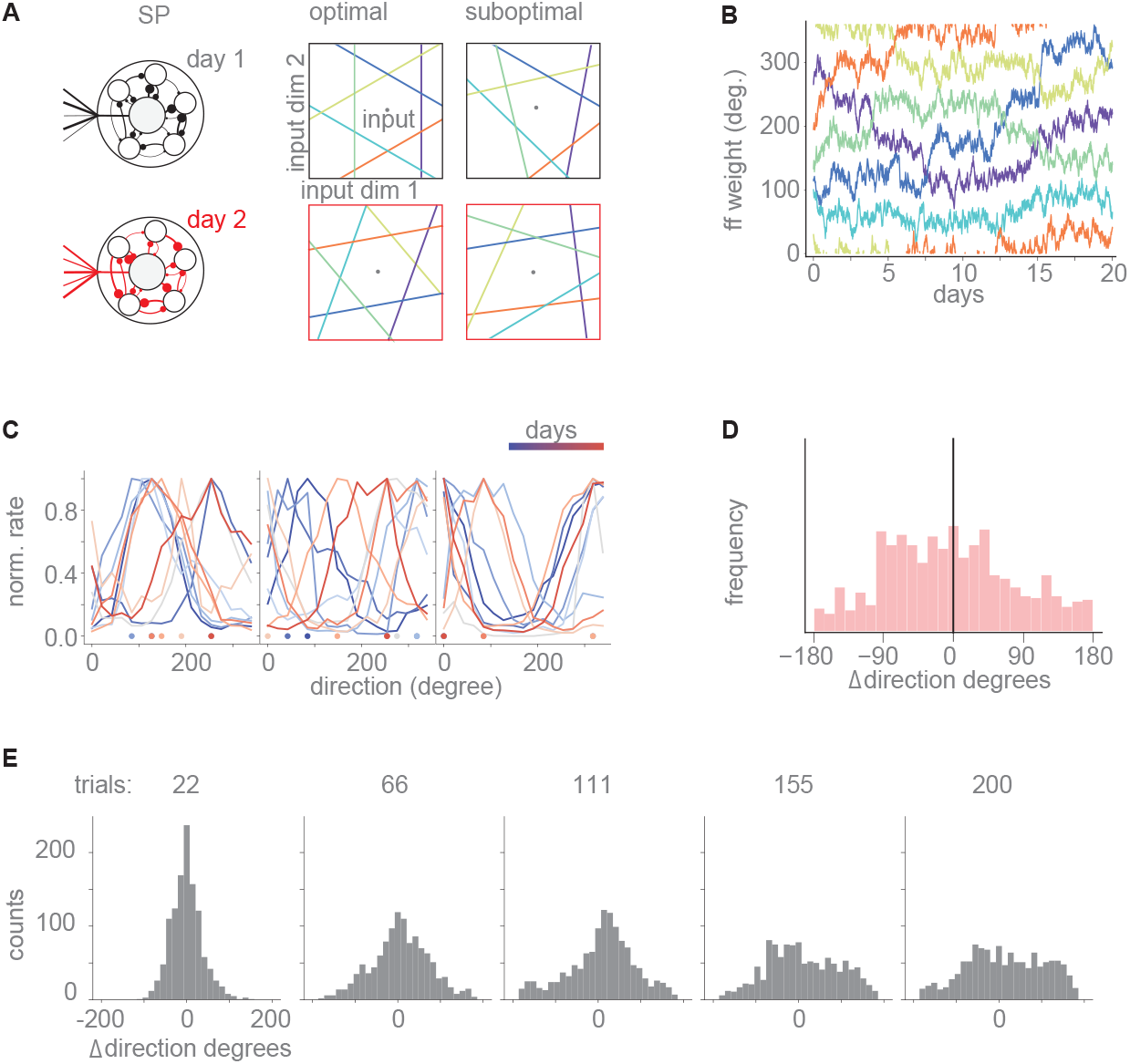
Synaptic weight changes lead to arbitrary tuning changes. A) Left: Illustration of synaptic weight changes across time and days. Middle: Individual neurons can converge to different coding angles but overall solution remains identical and optimal. Right: If noise is added to synaptic weight updates, neurons coding angles drift dynamically, resulting in suboptimal solutions. B) Feedforward weights of neurons (in angular degree) drift over time. This drift corresponds to random rotations of the bounding box in A. C) The tuning curves of three example neurons change position and shape over days as the coding angles drift. D) The distribution of preferred direction shifts span the entire spectrum from *−*180 to +180 degrees. (Notice that if this was orientation tuning and not direction tuning, the tuning changes would distribute between *±*90 instead.) E) The shape of preferred direction distribution shifts across trials.

